# Wnt1-Cre mediated deletion of BMP7 suggests a role for neural crest-derived BMP7 in retina development and function

**DOI:** 10.1101/2021.11.03.466838

**Authors:** Tiffany FC Kung, Pranidhi Baddam, Ruocun Liu, Devi Priyanka Maripuri, Ioannis S Dimopoulos, Ian M MacDonald, Yves Sauve, Daniel Graf

## Abstract

Neural crest (NC) contributes to various structures of the eye including cornea, ciliary body and retina. The association of NC-derived cells with hyaloid vessels in the form of pericytes is established. Similarly, persistence of NC-derived cells in the inner retina layer of the mature retina has been suggested. To date, no specific function has been attributed to them. NC-derived Bone morphogenetic protein 7 (BMP7) controls neurogenic properties in the brain and regulates glia differentiation. Here, we assessed the role of NC-derived BMP7 in the adult retina.

BMP7 expression was determined using Bmp7LacZ reporter mice. *BMP7* was expressed in GCL, IPL, OPL, and photoreceptors in P0, P14 and P30 retinas. Lineage tracing confirmed the presence of NC-derived cells in the GCL, INL, and ONL. Some but not all cells associated with vasculature. To test the function of NC-derived Bmp7, Bmp7^fl/fl^Wnt1cre (Bmp7^ncko^) mice were assessed by histological and functional methods. Loss of NC-derived cells in the GCL and INL and mild structural abnormalities were observed in the Bmp7^ncko^ retina. Electroretinography revealed reduced a wave under photopic conditions and b wave under both scotopic and photopic conditions. The neuronal circuitry in the inner retina appeared affected, evidenced by decreased Calbindin in the GCL, IPL and INL. In the outer retina, S-opsin was increased. BMP7 expression in the mutant retina was strongly decreased at birth, but increased expression from cells other than NC was observed in the adult retina. This was associated with an increase in IBA1, suggestive that loss of NC-derived BMP7 predisposes to development of gliosis-like changes in the adult retina. Overall, our data reveal an important contribution of NC-derived BMP7 for the development and function of the inner and outer retina.

## 1. Introduction

Derivatives of neural crest cells (NCCs) contribute to the formation of various structures in the adult eye, most notably the corneal endothelium and stroma of cornea, iris, and ciliary body (Gage et al., 2005; Whikehart, 2010; Williams and Bohnsack, 2015). Eye defects resulting from improper migration or differentiation of NCCs have mostly been associated with anterior segment dysgenesis syndromes and congenital glaucoma (Langenberg et al., 2008; Williams and Bohnsack, 2015). NCCs also contribute to the posterior segment, in particular pericytes of the transient choroidal hyaloid vasculature (Etchevers et al., 2001). Some NCC-derived cells persist in the adult retina as pericytes and vascular smooth muscle cells in the ganglion cell layer (GCL), the inner plexiform layer (IPL), and inner nuclear layer (INL) during postnatal stages (Trost et al., 2013). Furthermore Liu and colleagues have suggested that some lineage traced NCCs express markers for Müller glia, raising the possibility that a subset of Müller glia are NC-derived (Liu et al., 2014). Currently, no specific function has been described for these NC-derived retinal cells.

The adult retina is derived from neuroepithelium and pigmented epithelial cells that sequentially differentiate into 7 different layers: GCL, IPL, INL, outer plexiform layer (OPL), outer nuclear layer (ONL), inner segments/outer segments (IS/OS) and retinal pigment epithelium (RPE). The ganglion, horizontal, amacrine, and cone photoreceptor cells are the first to develop and mature, followed by rod photoreceptors, bipolar cells and Müller glia (Bassett and Wallace, 2012; Jeon et al., 1998), the major glial cell type of the retina. The retina converts light information into electrical signals and transmits the information to the visual processing centers in the brain via the process of phototransduction. Light travels through the eye to the photoreceptors where a signalling cascade hyperpolarizes the photoreceptors. The photoreceptors synapse onto rod and cone bipolar cells, which then synapse onto the ganglion cells from where the electrical information travels to the brain (Masland, 2012). Within this process, Müller glia have several functions. Regularly spaced and spanning the entire thickness of the neuroretina, they are involved in regulating potassium ion flux, serve as structural support for neurons, and modulate neuronal and neurovascular coupling (Reichenbach and Bringmann, 2013). In contrast to mammals, zebrafish Müller glia maintain the life-long potential to regenerate all major retinal cell types (Hoang et al., 2020; Martins et al., 2020).

In the brain, the NC-derived meninges provide critical trophic support during embryonic neural development by regulating the neurogenic properties of radial glia cells (Segklia et al., 2012; Siegenthaler and Pleasure, 2011), the life-long neurogenesis in the dentate gyrus, and dentate neural stem cells (Choe et al., 2013). This process is in part mediated by BMP7, an important signalling molecule needed to maintain normal PAX6 expression, including after lens placode induction (Segklia et al., 2012; Wawersik et al., 1999). BMP signaling also controls Müller glia differentiation, and the exposure of retinal astrocytes and Müller glia to BMP7 results in reactive gliosis (Dharmarajan et al., 2014; Wawersik et al., 1999). *BMP7* mutations in humans are associated with various ocular abnormalities ranging from anophthalmia to microphthalmia (Wyatt et al., 2010). Complete loss of BMP7 in mice results in anophthalmia (Zouvelou et al., 2009), highlighting its significance for eye development. Despite this, little is known about the requirement of BMP7 for the adult retina. No genetic studies using tissue-specific deletion of BMP7 in the retina have been performed. Given that NC-derived BMP7 is important for neural development and stem cell maintenance, and a subset of Müller glia might be NC -derived, we asked whether BMP7 is expressed in the adult retina and whether NC-specific BMP7 deletion would affect retina development and function.

In this study, we demonstrate that BMP7 remains expressed in various compartments of the adult eye, including the retina. Loss of BMP7 in NCC (Bmp7^ncko^) results in loss of NCC in the GCL and INL, discrete cell organization defects in the outer retina, and abnormalities to neuronal circuitry in the inner retina. NC-specific BMP7 loss leads to increased expression of BMP7 by non-NC-derived expression in the adult retina. Correlating cellular and molecular changes in the mutant retina to changes in light perception and signalling using electroretinogram (ERG), we identified a requirement of NC-derived BMP7 for normal vision. To our knowledge, this is the first description of how an NC-derived growth factor ensures normal function of the adult retina. Our data not only demonstrate how subtle changes to cellular and molecular properties translate to visual defects in the adult eye, but also offer possible new avenues for exploiting the plasticity of the retina to potentially correct retinal dysfunction.

## 2. Methods

### 2.1. Ethics Statement

All procedures were conducted in accordance with the Canadian Council on Animal Care guidelines and approved by the Animal Care and Use Committee of the University of Alberta (protocol: AUP1149).

### 2.2. Animals

Bmp7LacZ reporter mice (Godin et al., 1998), BMP7 neural crest knock-out mice (Bmp7^fl/fl:Wnt1cre^, subsequently referred to as Bmp7^ncko^ or mutant mice) were generated as previously described (Malik et al., 2020; Zouvelou et al., 2009). Mouse lines were kept on a C57BL/6J background. Bmp7^fl/fl^ mice served as controls (Bmp7^ctrl^), except for lineage tracing, where Bmp7^wt:Wnt1cre^ mice were used. Lineage tracing was done using mT/G mice (*Gt(ROSA)26Sor*^*tm4(ACTB-tdTomato,-EGFP)Luo*^/J) (Muzumdar et al., 2007). Due to the exploratory nature of this study, *a priori* power was not calculated, but post-hoc power and effect sizes are provided. Male and female mice were used for all experiments. For histological/immunofluorescence analysis, a minimum of three biological replicates were analyzed.

### 2.3. LacZ Staining

LacZ staining of Bmp7^Wt:LacZ^ enucleated mouse eyes were performed as previously described (n=3/age) (Malik et al., 2020). LacZ stained eyes were fixed using Davidson’s fixative solution (Richmond, n.d.) as described below.

### 2.4. Tissue Preparation and Histology

Animals were euthanized, and enucleated eyes were fixed with Davidson’s fixative solution (Richmond, n.d.) for 12 hours, stored in 50% ethanol, and processed for paraffin embedding and sectioning, as previously described (Baddam et al., 2020). Eyes were sectioned in the sagittal orientation at a thickness of 7μm. Sections 70 to 140 μm ventral to the optic nerve were used for staining. Hematoxylin and eosin staining was performed as previously described (Ellis, n.d.; Malik et al., 2020)

### 2.5. Immunofluorescence

Molecular markers were investigated in Bmp7^ctrl^ and Bmp7^ncko^ mice (n=3) and lineage tracing experiments were conducted in Bmp7^ctrl^, Bmp7^ncko^, and Bmp7^Wt:Wnt1cre^ mice (n≥3). Immunofluorescence and imaging were done as previously described using paraffin sections (Malik et al., 2020). Sections were deparaffinized. Antigen retrieval was performed using 10mM sodium citrate buffer in a microwave. Sections were blocked with the appropriate goat or donkey serum in Tris-buffered saline (TBS). Primary antibodies were incubated at 4°C overnight. Primary antibodies used were: Calbindin (Calb; Santa Cruz, sc28285), Neurofilament Heavy (NF-H; Abcam, ab187374), Glial Fibrillary Acidic Protein (Gfap; Abcam, ab4674), Pax6 (prb-278P), Rhodopsin (Rho; Abcam, ab98887), Recoverin (Rcvrn; Abcam, ab5585), Blue Opsin (S-op; short-wavelength; Millipore Sigma, ab5407), Red & Green Opsin (R&G-op; medium wavelength; Millipore Sigma, ab5405), Tomato Lectin (Invitrogen L32470 DyLight 488), Bone Morphogenetic Protein 7 (Bmp7; Abcam, ab84684) and Green Fluorescent Protein (Gfp; Abcam, ab6556, provided by Luc Berthiaume, Department of Cell Biology, University of Alberta). Secondary antibodies were: donkey anti-rabbit Alexa Flour (AF)-647 (6440-31), goat anti-mouse IgG1AF-647 (A21240), goat anti-mouse IgG2a AF-647 (A21241), goat anti-mouse IgG3 AF-594 (A21155), and goat anti-chicken AF-594 (ab150172), donkey anti-chicken AF-488 (AB_2340375). Slides were mounted using 2.5% DABCO Mowiol and stored in the dark to avoid bleaching, at room temperature. All images were captured on an Olympus IX73 microscope at 20x magnification, except for S-op and R&G-op which was also imaged at 10x. These images were compiled and overlaid using FireAlpaca (v. 2.2.10). Photoreceptors were counted from three independent biological replicates. Whole retinal sections were used to count photoreceptors positive for S-op (Blue cones) and R&G-op (Red and Green cones). A single rater counted the photoreceptors and was blinded to the genotype of the mouse. Image intensities were normalized to DAPI staining, and true staining was confirmed through comparison to antibody negative controls and auto-fluorescent controls. All slides were incubated in antibodies for the same length of time, and were imaged at the same parameters (i.e., intensity, gain, exposure).

### 2.6. Optical Coherence Tomography (OCT)

One-month-old (P30) Bmp7^ctrl^, Bmp7^ncko^ (n=4/genotype) were anesthetized by intraperitoneal injection of 150mg/kg ketamine /10mg/kg xylazine; phenylephrine and tropicamide were applied to dilate the pupil and prevent dryness. Mice were placed into a stereotactic rotational cassette and the retina was scanned using the Bioptigen Envisu spectral domain ophthalmic imaging OCT system. Data analysis was done using InVivoVue Clinic imaging software. To measure retinal thickness, calipers were drawn across all retinal layers (RNFL: retina nerve fiber layer; GCL: ganglion cell layer; IPL: inner plexiform layer; INL: inner nuclear layer; OPL: outer plexiform layer; ONL: outer nuclear layer; IS/OS: inner segments/outer segments; RPE: retina pigmented epithelium), for both the superior and inferior retina. Each mouse (2 eyes per mouse) was evaluated by two researchers with high intra- (kappa: 0.94) and inter-rater reliability (ICC: 0.83) blinded to mouse genotype at the time of assessment. Due to poor image quality of the inferior retina, one Bmp7^ncko^ mouse was excluded from all analysis for the inferior retina.

### 2.7. Electroretinogram (ERG)

Mice were anesthetized by intra-peritoneal injection of 150mg/kg ketamine /10mg/kg xylazine. Scotopic conditions were tested first, followed by photopic conditions using 10 μs light flashes (0.3-300Hz bandpass without notch filtering). Scotopic flashes ranged from -5.22 to 2.86 log cd s/m^2^, and photopic flashes ranged from -1.6 to 2.9 log cd s/m^2^ in 30 cd/m^2^ background illumination. Body temperature of the mice was maintained at 38^º^C. Dark (scotopic) and light (photopic)-adapted electroretinography (ERG) was performed as previously described (Cheng et al., 2020) on a total of nineteen P30 mice per group (N=38) using the Espion E 2 system (Diagnosys, LLC, Littleton, MA). For scotopic conditions, mice were dark-adapted for 2 hours prior to measurement and handled under dim red light. Researchers were blinded for mouse genotype during measurement and analysis.

### 2.8. Mining of scRNA Seq data

The scRNA seq expression matrix of mouse retinal progenitor cells across 10 time-points of retinal development (GEO Accession ID GSE118614, along with its metadata) (Clark et al., 2019) was processed using Seurat to normalize data allowing comparison between cells using Log Normalization, detection of genes with variable expression patterns, and clustering and identification of cluster-specific marker genes to annotate the cells with cell types. Results were visualized using non-linear embedding methods including tSNE and UMAP, and a sparse matrix R data file (.rds) file was generated. Sceasy was used to convert the file format from a Seurat’s .rds object to an annData’s .h5ad object, which was then loaded into an interactive single cell transcriptome data explorer, cellxgene. The processed data was then loaded into the visualizer using the command: cellxgene launch retina_ids_11_final.h5ad --open. The link to the processed data file is available for access using https://github.com/PriyankaMaripuri/Retina.

### 2.9. Statistics

*A priori* power was not calculated as this was an exploratory analysis. Post-hoc power analyses are provided for all significant results. The Shapiro-Wilk test for normality was used to determine normality. Photoreceptor counts, OCT retinal layer lengths, and ERG comparisons were assessed using unpaired t-tests. For ERG analyses, intensities below the detection limit (0 amplitude response) were omitted from analyses. All data are presented as mean ± SD.

## 3. Results

### 3.1. BMP7 expression is dynamic and overlaps with NCCs in the post-natal eye

To understand when and where BMP7 is expressed in the post-natal eye, its expression was mapped using Bmp7LacZ reporter mice. At birth (Post-natal day 0, P0), no expression was observed in the cornea (Figure 1A), but clear expression was seen in the ciliary body (Figure 1B) and the RPE of the neural retina (Figure 1C).

**Figure 1.**
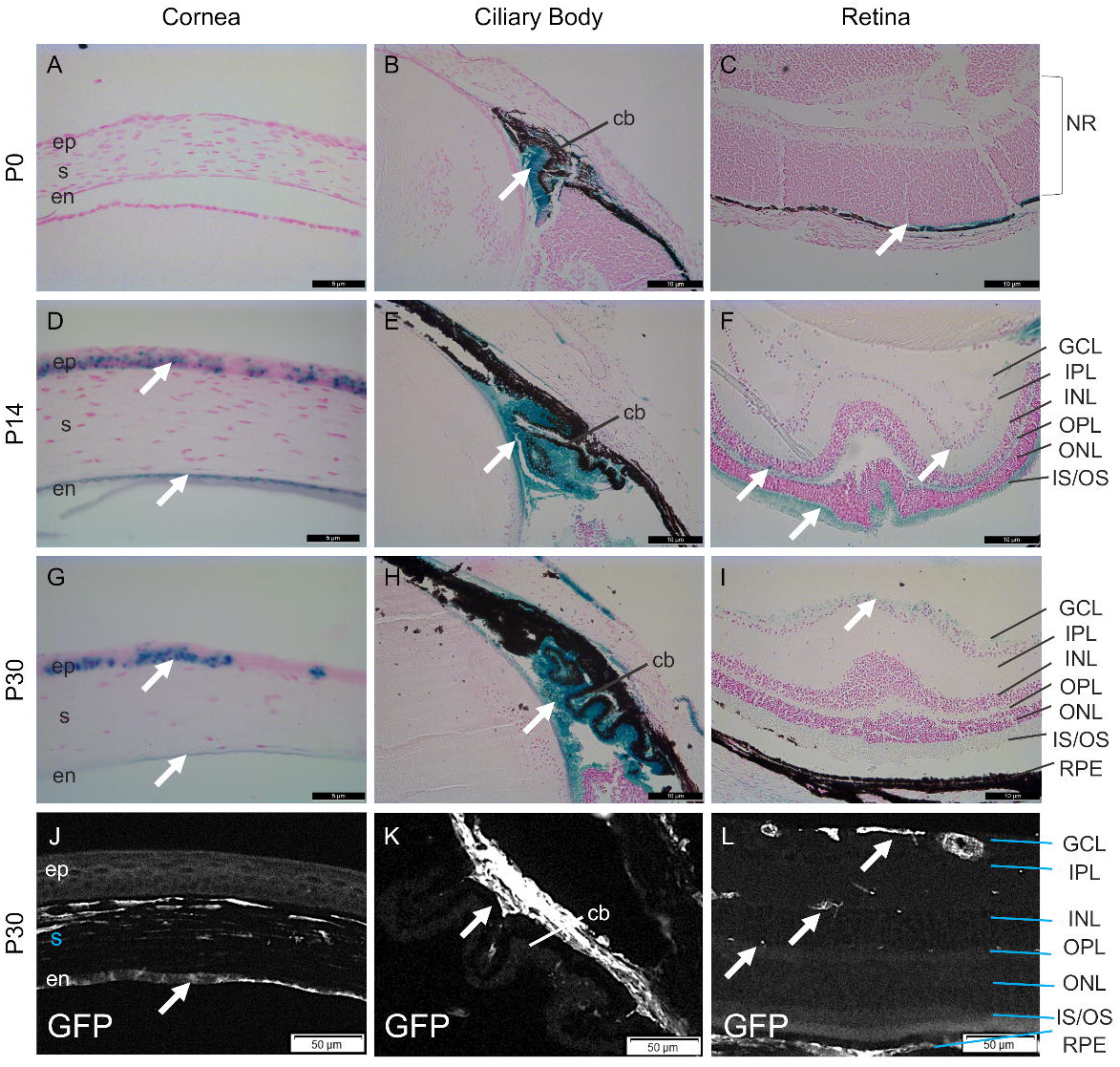
Dynamic expression of BMP7 and NCC in the mouse eye. Sagittal sections of paraffin-embedded eyes counterstained with nuclear fast red from Bmp7LacZ reporter mice demonstrate minimal expression of BMP7 (blue) in the cornea **(A)**, ciliary body **(B)** and retina **(C)** at P0. At P14, BMP7 is expressed in the epithelium and endothelium of the cornea **(D)**, the cells lining the lens and the ciliary body **(E)** and the GCL, OPL and IS/OS of the retina **(F)**. Expression is sustained only in the cornea epithelium **(G)**, ciliary body **(H)** and GCL of the retina **(I)** at P30. GFP immunofluorescence staining(McCabe and Berthiaume, 1999) (white) for NCC using Bmp7^Wt:Wnt1cre^ eyes revealed the presence of NCC in the stroma and endothelium of the cornea **(J)**, ciliary body **(K)** and the GCL and INL of the retina **(L)** at P30. P0: postnatal day 0; P14: postnatal day 14; P30: postnatal day 30; ep: epithelium; s: stroma; en: endothelium; cb: ciliary body; nr: neural retina; gcl: ganglion cell layer; ipl: inner plexiform layer; inl: inner nuclear layer; opl: outer plexiform layer; onl: outer nuclear layer; is/os: inner segments/outer segments; rpe: retinal pigment epithelium; GFP: green fluorescent protein.

At P14, when eyes open and rapid eye growth is observed in mice (Shen and Colonnese, 2016; Tkatchenko et al., 2010), both the corneal epithelium and endothelium expressed BMP7 (Figure 1D). Expression in the growing ciliary body persisted and appeared to extend anterior (towards to the iris) and posterior (towards the developing GCL of the retina) (Figure 1E). Within the retina, BMP7 was strongest in the IS/OS layer, but expression extending to the OPL and GCL was apparent (Figure 1F).

At P30, BMP7 expression in the corneal epithelium became more restricted into clusters, while expression in the endothelium appeared comparatively weaker (Figure 1G). In the ciliary bodies, expression remained ubiquitous (Figure 1H), whereas in the retina BMP7 expression was limited to the GCL (Figure 1I). High magnification images of Bmp7 expression demonstrated in Figure S1.

To assess which of these BMP7 expression domains overlapped with neural crest-derived cells and structures, lineage tracing was performed using P30 Bmp7^Wt:Wnt1cre^ retinas (n=3 mice). NCC-derivatives could be observed in stroma and endothelium of the cornea (Figure 1J), the pigmented cells of the iris and within the ciliary bodies extending towards the retinal pigmented epithelium (Figure 1K). Within the retina, lineage-traced cells were apparent in the GCL, IPL, INL and OPL (Figure 1L). A negative control for NCC staining is demonstrated in Figure S2. The partial overlap between lineage tracing and BMP7 expression prompted us to assess if NC-derived BMP7 is important for postnatal eye development and function.

### 3.2. Bmp7^ncko^ mice display molecular abnormalities, but no structural differences

Gross morphological analysis revealed that eyes of Bmp7^ncko^ mice were of comparable size. No apparent alterations to the cornea were apparent, but we noted that pupils were slightly smaller, although not significant, after pharmacological dilation (Figure S3). To gauge if there were any obvious structural defects in the retina, we characterized the retina using OCT and histology. Whereas no statistically significant changes to the retinal layers was observed at either inferior (p≥0.06) or superior (p≥0.13) positions (Figures 2A-D), histological assessment identified some mild disorganization of neuronal cell bodies in the ONL suggesting disrupted photoreceptor stacking when compared to controls (Figure 2E-F). A reduction or loss of NC-derived cells was observed specifically in the GCL and INL of Bmp7^ncko^ retinas compared to Bmp7^Wt:Wnt1cre^ mice, whereas NC-derived cells at the OPL appeared unaffected. Loss of BMP7 did not affect cellularity of NC-derived components lining the RPE (Figure 2G-H). We thus asked next if loss of BMP7 resulted in functional impairments associated with inner and outer retina.

**Figure 2.**
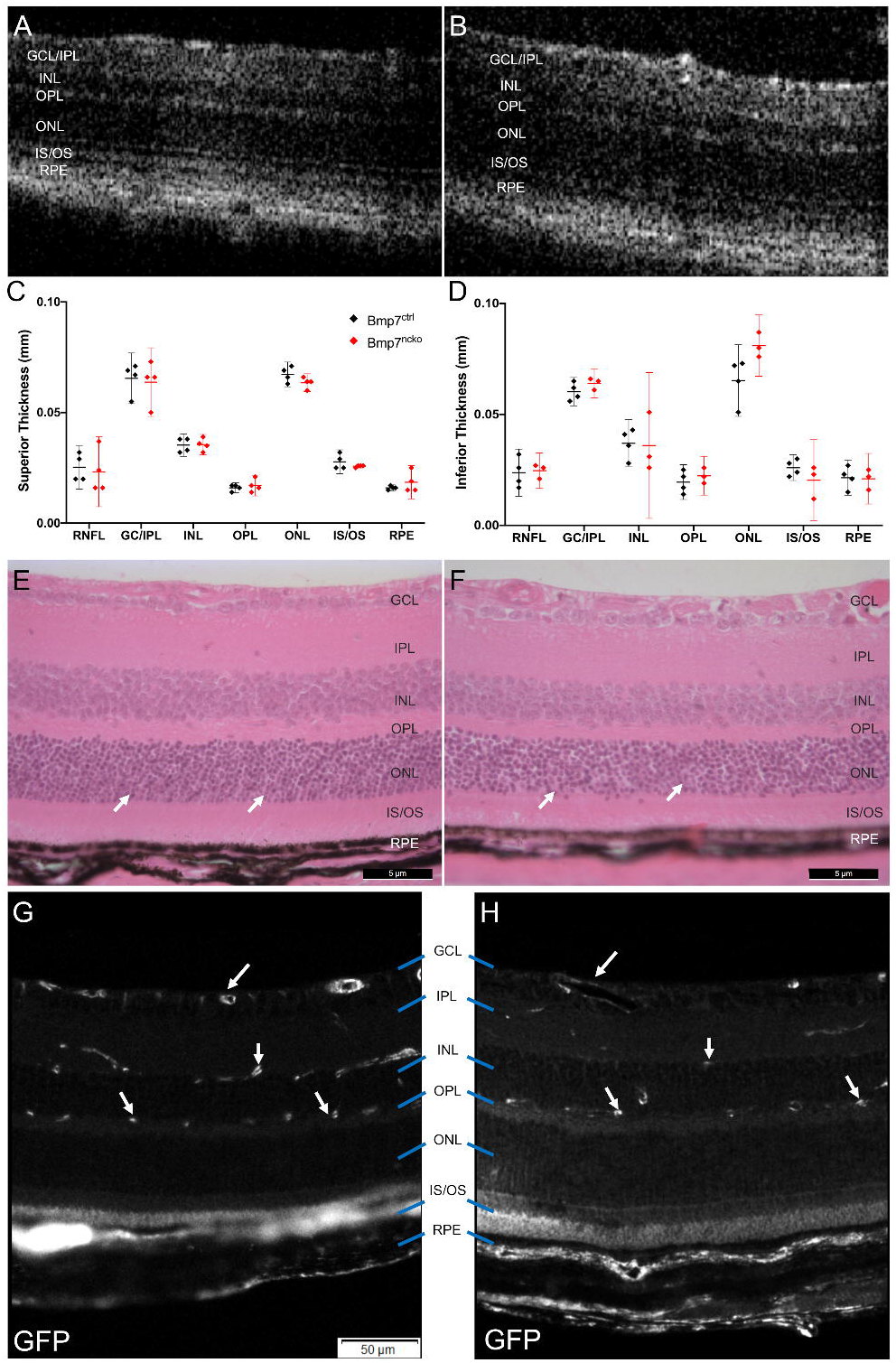
Bmp7^ncko^ mice demonstrate abnormal cell organization in ONL and loss of NCC in the inner retina with no obvious changes to thickness of retina layers. **Retinal layer** thickness assessment using OCT imaging of P30 Bmp7^ctrl^ **(A)** and Bmp7^ncko^ **(B)** mice (n=4/genotype) showed no differences to retinal layer thickness at both superior **(C)** and inferior **(D)** locations. Mildly disorganized cell stacking (arrows) in the ONL was observed in Hematoxylin and Eosin stained P30 Bmp7^ncko^ **(F)** retina compared to Bmp7^ctrl^ **(E)**. Loss of GFP-positive NCC (white) was observed in the GCL and INL (arrows) of P30 Bmp7^ncko^ **(H)** mice in comparison to Bmp7^Wt:Wnt1cre^ **(G)** mice. OCT: Optical coherence tomography; P30: postnatal day 30; rnfl: retina nerve fiber layer; gcl: ganglion cell layer; ipl: inner plexiform layer; inl: inner nuclear layer; opl: outer plexiform layer; onl: outer nuclear layer; is/os: inner segments/outer segments; rpe: retinal pigment epithelium; GFP: green fluorescent protein.

### 3.3. Bmp7^ncko^ mice exhibit inner retina deficits

ERG assessments performed on P30 Bmp7^ctrl^ (black) and Bmp7^ncko^ (red) mice (n=19/genotype) revealed significant reductions in the a-wave (indicator of outer retina function (Creel, 1995)) under scotopic conditions (Figure 3A; p=0.04), but not under photopic conditions (Figure 3B; p=0.22, Cohen’s d= 4.79). In contrast, the amplitudes of the b-wave (an indicator of inner retina function (Creel, 1995)) under both scotopic (Figure 3C; p=0.0009, Cohen’s d=8.8 with 100% power) and photopic conditions (Figure 3D; p=0.008, Cohen’s d= 8.49 with 100% power) were reduced. The reduced ratio of scotopic b-wave/a-wave pointed towards a more pronounced inner retina defect (Figure 3E; p=0.002, d=1.72 with 99% power). The flicker response, which identifies the status of cone photoreceptor function (Verma and Pianta, 2009), also showed a significant reduction (Figure 3F; p=0.05, d= 7.5 with 100% power) reflecting decreased function of cone photoreceptors. The implicit time, a measure of the time it takes to generate a- or b- wave signals (Creel, 1995), was not significantly different for the a-wave under both scotopic and photopic conditions (Figure 3G-H; pε0.70), suggestive of normal phototransduction. The implicit time for the scotopic b-wave was significantly shorter (Figure 3I; p=0.01, d=1.52 with 99% power), while the photopic b-wave implicit time showed no significant difference (Figure 3J; p=0.91). The selectivity and severity of the various visual deficits observed by ERG were surprising and could not easily be explained by a mildly disorganized ONL. We thus sought to further characterize the molecular and cellular composition of the inner and outer retina using selected antigen markers by immunofluorescence.

**Figure 3.**
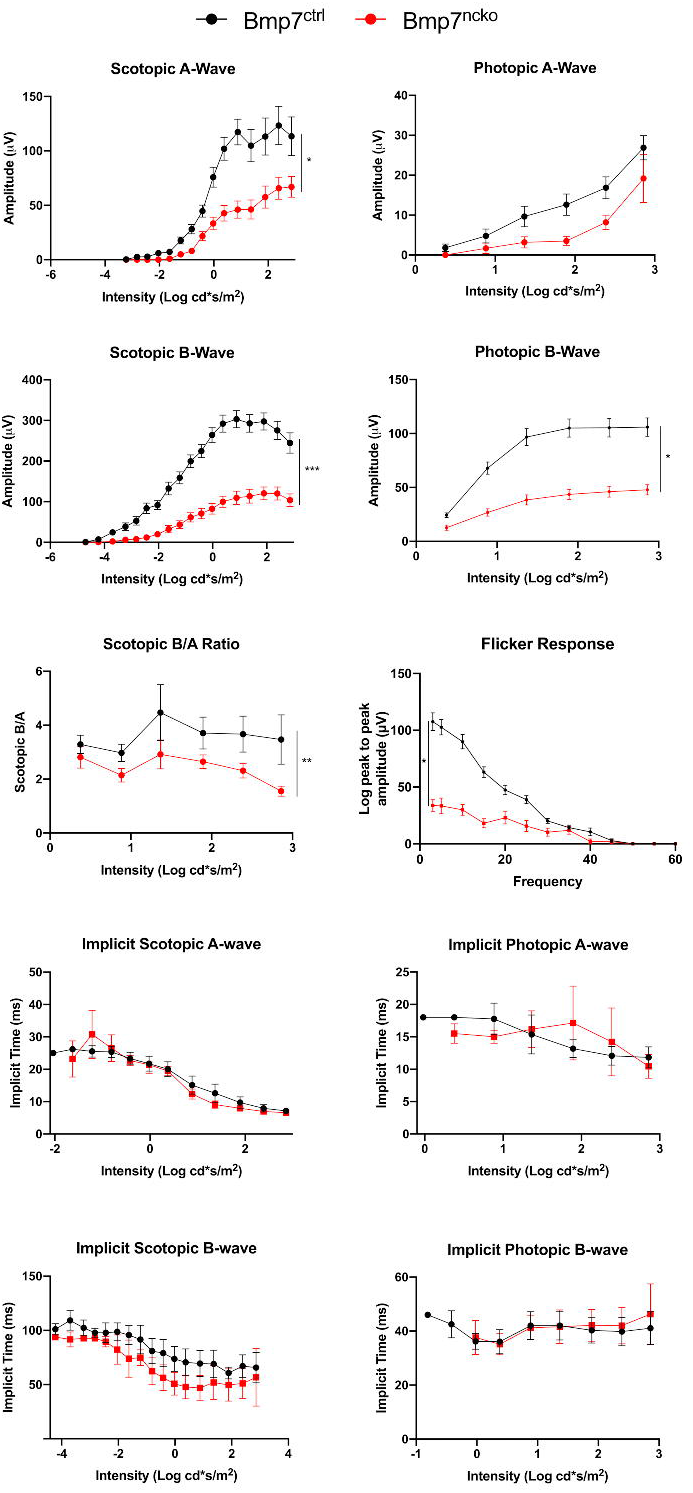
Bmp7^ncko^ mice have a more pronounced inner retina functional deficit than outer retina. Electroretinogram of P30 mice (n=19/genotype) indicated a significant reduction in a-wave (outer retina function) of Bmp7^ncko^ mice in scotopic **(A)** but not photopic **(B)** conditions. Assessment of the b-wave (inner retina function) also demonstrated a significant reduction under both scotopic **(C)** and photopic **(D)** conditions. A reduction in the scotopic b/a ratio **(E)** and a delay in flicker response **(F)** of Bmp7^ncko^ mice retina confirm the severity of the inner retina defect. The implicit time of a-wave under both scotopic **(G)** and photopic **(H)** conditions demonstrated no significant differences. However, a significant reduction in the implicit time of scotopic b-wave **(I)** was observed, with no changes to implicit time of photopic b-wave **(J)**. For all statistical analyses, intensities that were too low for rats of both groups (0 amplitude response) were omitted. Black: Bmp7^ctrl^; Red: Bmp7^ncko^. P30: postnatal day 30.

### 3.4. Abnormal neuronal circuitry in the inner retina of P30 Bmp7^ncko^ mice

As visual deficits in mutant mice were primarily associated with inner retina function, we assessed first the neuronal organization of the retina. DAPI stain is shown for reference (Fig. 4A-B). Calbindin (CALB), a calcium-binding protein broadly expressed in bipolar, amacrine, ganglion, and cone photoreceptor cells (Gu et al., 2016; Morona et al., 2008) was reduced in the GCL and INL of mutant mice (Figure 4C, D). Heavy chain neurofilament (NF-H), which stains retinal ganglion cells to assess neuronal damage (Kashiwagi et al., 2003), was increased in the GCL (Figure 4E-F). Glial fibrillary acidic protein (GFAP), which stains glial cells and astrocytes (Sarthy et al., 1991), revealed potential differences to cell composition in IPL and OPL in the mutant (Figure 4G-H, G’-H’). Ionized calcium binding adaptor molecule 1 (IBA1), a microglia maker(Tassoni et al., 2015), was increased in GCL, INL and ONL in Bmp7^ncko^ mice. Paired box protein 6 (PAX6), which is largely confined to retinal ganglion cells and amacrine cells at this age (Figure 4I-J) (Li et al., 2007; Riesenberg et al., 2009), was markedly reduced in the GCL and INL (Figure 4I-J). The disruption of normal CALB, NF-H and PAX6 expression in the GCL and INL, as well as the differences in glial cell organization, suggest atypical neuronal circuitry of the inner retina.

**Figure 4.**
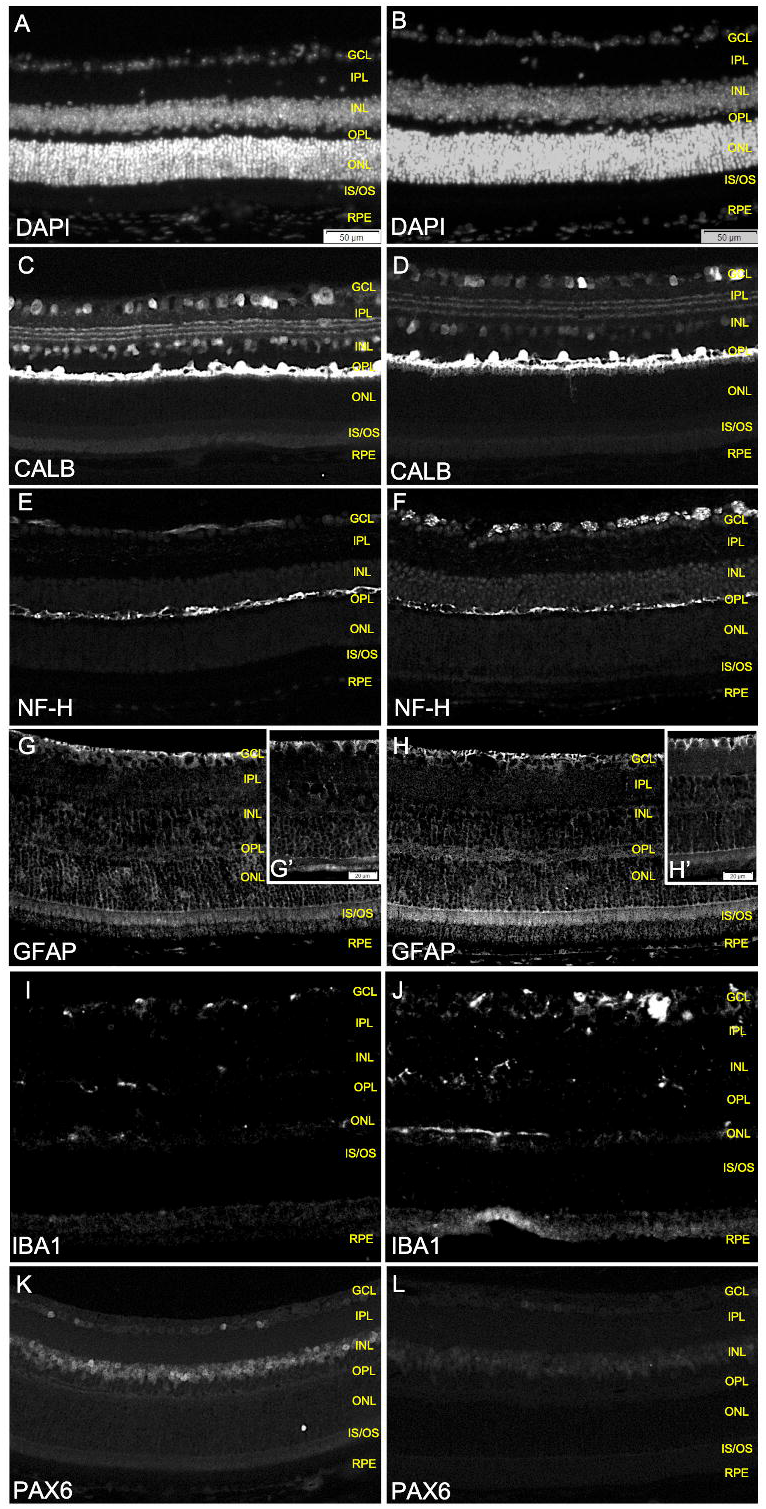
Abnormal neuronal organization in the inner retina of Bmp7^ncko^ mice. Bmp7^ctrl^ **(A, C, E, G, G’, I, K)** and Bmp7^ncko^ **(B, D, F, H, H’, J, L)** retina were stained for various neuronal proteins and DAPI for nuclei **(A-B)** using immunofluorescence. Reduced expression of CALB **(C-D)** was observed in GCL and INL of the mutant retina. Increased expression of NF-H **(E-F)** was observed in the GCL of Bmp7^ncko^. A slight increase of GFAP **(G-H)** expression was observed in the GCL. **(G’-H’)** Higher magnification images of the GFAP expression in the GCL. Expression of IBA1 **(I-J)** was increased in GCL, INL and ONL layers of Bmp7^ncko^. Expression of PAX6 **(K-L)** was also reduced in the GCL and INL of the mutant mice. Immunostaining was performed on paraffin-embedded P30 retina (n=3/genotype). CALB: calbindin; NF-H: neurofilament heavy; GFAP: glial fibrillary acidic protein; IBA1: Ionized calcium binding adaptor molecule 1; PAX6: paired box protein 6. gcl: ganglion cell layer; ipl: inner plexiform layer; inl: inner nuclear layer; opl: outer plexiform layer; onl: outer nuclear layer; is/os: inner segments/outer segments; rpe: retinal pigment epithelium. P30: postnatal day 30.

### 3.5. Impaired organization of cone photoreceptors in the outer retina of Bmp7^ncko^ mice

Next, we assessed potential molecular changes that would correlate with the a-wave findings. DAPI stain is shown for reference (Fig. 5A-B). Rhodopsin (RHO) (Figure 5C-D) and Recoverin (RCVRN) (Figure 5E-F) are predominantly expressed in rod photoreceptors that govern vision in dim light (scotopic conditions). RHO was reduced in the IS/OS layer, while RCVRN was reduced in both the ONL and IS/OS layers, indicating an effect of NC-derived BMP7 on photoreceptors. Wavelength-specific cone photoreceptors that govern colour vision under bright light (photopic conditions) also showed changes. Blue opsin (S-OP) confined to the ventral aspect of the retina (Nadal-Nicolás et al., 2020; Ortín-Martínez et al., 2014) showed an increase in the IS/OS layer of the Bmp7^ncko^ retina (Figure 5G, H). Red and Green opsin (R&G-OP) showed no apparent numerical changes (Figure 5I, J). Quantification confirmed a significant increase of S-OP in Bmp7^ncko^ retina (Figure 5K; p=0.01, d=3.71 with 92% power), with no changes to the number of R&G-opsin expressing cones (Figure 5L; p=0.14). The cumulative deficits to neuronal circuitry in the inner retina and rod/cone photoreceptor abnormalities in the outer retina indicate that NCC-derived BMP7 exerts effects on multiple cell types in the adult retina. However, it is still unclear which retinal cells are NC-derived and when NC-derived BMP7 is required.

**Figure 5.**
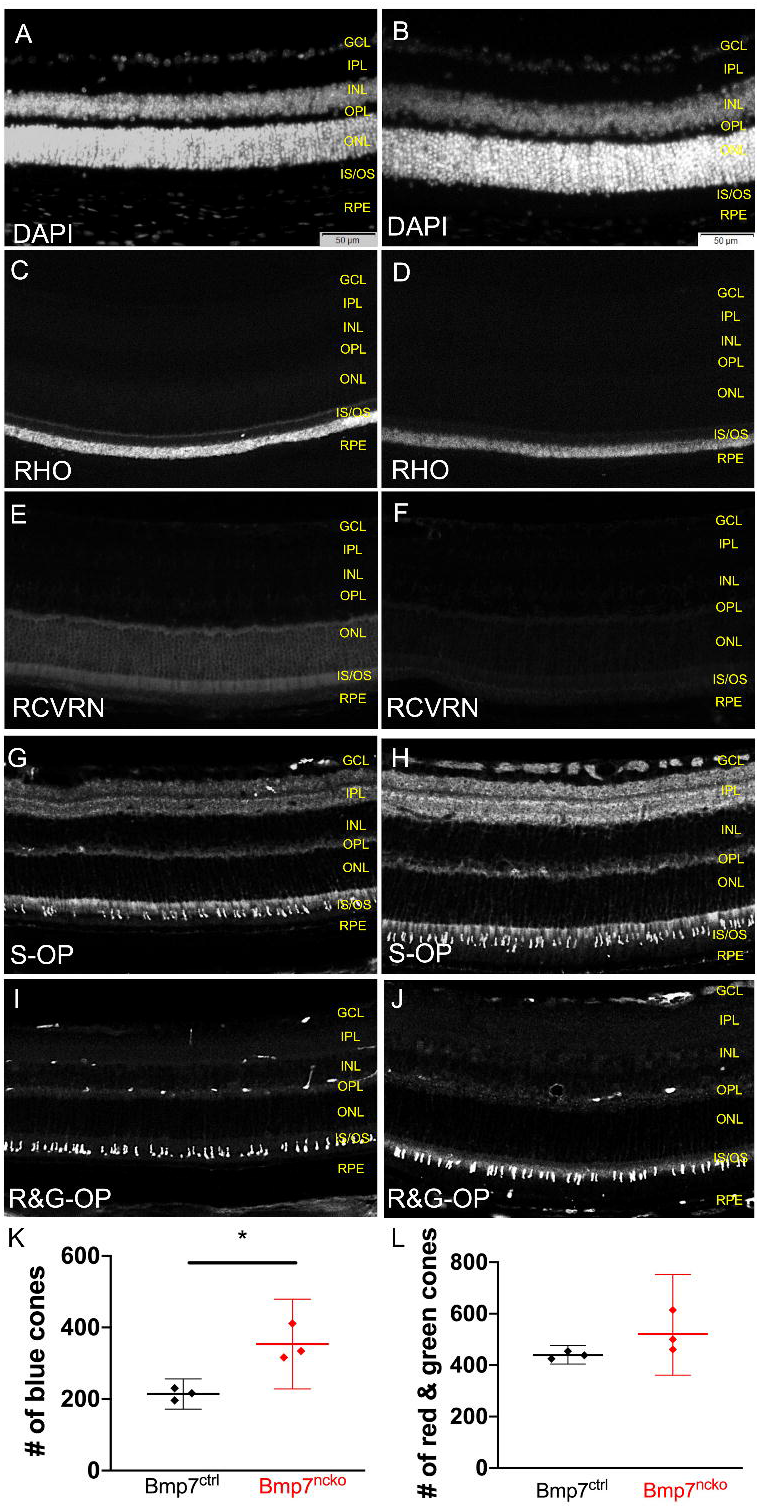
Increase in blue-opsin expressing cone photoreceptors in the outer retina of Bmp7^ncko^ mice. P30 Bmp7^ctrl^ **(A, C, E, G, I)** and Bmp7^ncko^ **(B, D, F, H, J)** retina (n=3/genotype) were immune-stained for various proteins expressed in rod and cone photoreceptor and DAPI for nuclei **(A-B)** using immunofluorescence. Staining for rod photoreceptors using RHO **(C-D)** and RCVRN **(E-F)** demonstrated a reduction in IS/OS layer and ONL of the BMP7^ncko^ mice. Staining for cone photoreceptors using S-OP **(G-H)** and R&G-OP **(I-J)** demonstrated an increase in blue cones and no significant changes to red and green cones. Cone photoreceptor quantification indicated an increase in blue-opsin expressing cones **(K)** with no difference in red and green opsin-expressing cones **(L)** in Bmp7^ncko^ mice. Immunostaining was performed on paraffin-embedded P30 retina (n=3/genotype). RHO: rhodopsin; RCVRN: recoverin; S-OP: blue opsin; R&G-OP: red and green opsin. gcl: ganglion cell layer; ipl: inner plexiform layer; inl: inner nuclear layer; opl: outer plexiform layer; onl: outer nuclear layer; is/os: inner segments/outer segments; rpe: retinal pigment epithelium. P30: postnatal day 30.

### 3.6. Persistent neural crest cells associate with cells in the INL and ONL in addition to the vasculature

Earlier studies established that some NC-derived cells persist in the inner nuclear and ganglionic cell layers of the retina as vasculature-associated pericytes to at least 6-weeks of age (Trost et al., 2013). To test if all NCCs associate with vasculature pericytes or whether some cells contribute to other cell types, we colocalized NCCs (GFP lineage tracing) with vasculature (Tomato Lectin (TL)), neurons (NF-H) and glial cells (GFAP). Colocalization with GFP and TL in P0, P14 and P30 Bmp7^Wt:Wnt1cre^ mice revealed that NCCs were frequently associated with vasculature (Figure 6A) but not all vasculature associated with NCCs. We also observed cells, predominantly at P0 and P14 that did not associate with vasculature, as evident by absent TL staining (Figure 6A white arrows). We also occasionally observed a striped pattern of GFP positive cells in 1 of 3 Bmp7^Wt:wnt1cre^ mice at all three ages investigated. Higher magnification images for Figure 6A as shown in Figure S4.

**Figure 6.**
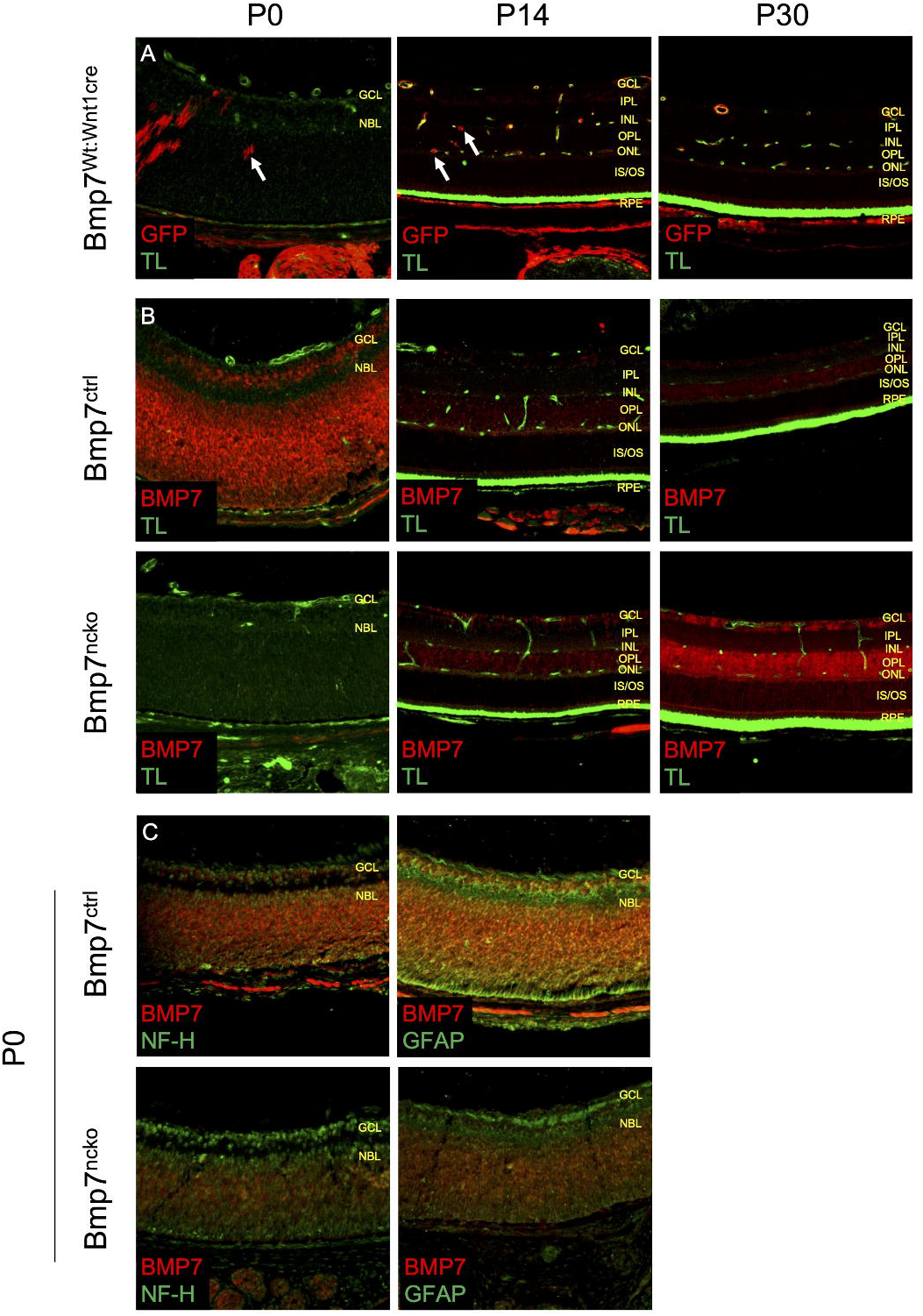
Persistent neural crest cells associate with cells in the INL and ONL and deletion of BMP7 increases BMP7 expression in the mature retina. GFP and BMP7 were co-localized with tomato lectin (TL; **A, B**), NF-H, and GFAP (**C**) to test if all NCCs associate with vasculature or other retina cell types. At P0 and P14, the majority of GFP-positive cells colocalize with TL but few GFP-positive cells in the INL and ONL are negative for TL (**A**, white arrows). A striped GFP staining pattern was observed in 1 of 3 Bmp7^Wt:wnt1cre^ mice at P0 and P30, resembling findings presented in Liu et al., 2014.^7^ Similarly, BMP7 expression in P0, P14, and P30 and Bmp7^ncko^ retina (**B**) revealed expression being confined mostly to the GCL and OPL layers, where TL staining is also visible. In Bmp7^ctrl^ mice, BMP7 expression decreased as the retina matures, however in Bmp7^ncko^ mice, a dramatic loss of BMP7 was observed initially at P0 with a gradual increase of expression evident at both P14 and P30 timepoints. Neuronal (NF-H) and glial (GFAP) cell markers, were colocalized with BMP7 at P0 (**C**) where a significant amount of BMP7 is associated with GFAP positive glial cells whereas no comparable colocalization was observed with NF-H. NF-H: neurofilament heavy; GFAP: glial fibrillary acidic protein; GFP: green fluorescent protein; gcl: ganglion cell layer; inl: inner nuclear layer; opl: outer plexiform layer; onl: outer nuclear layer rpe: retinal pigment epithelium. P0: postnatal day 0; P14: postnatal day 14; P30: postnatal day 30.

### 3.7. NCC-specific deletion of BMP7 increases BMP7 expression in the mature retina

Findings from Bmp7LacZ reporter mice (Figure 1) showed much broader BMP7 expression than could be attributed to NCCs. To test whether NC-specific BMP7 deletion results in changes of BMP7 expression in the retina, we probed for BMP7 expression in P0, P14, and P30 (n=2 at P30, n=1 at P0 and P14) Bmp7^ctrl^ and Bmp7^ncko^ retinas (Figure 6B). At P0, BMP7 was broadly expressed except at the interface between the GCL and NBL (neuroblastic layer). At P14 and P30, BMP7 expression appeared weaker and was restricted to the GCL, OPL and ONL. In P0 Bmp7^ncko^, BMP7 expression was consistently reduced (n=4) but expression was variable ranging from near absent to weak expression (compare Figure 6A, C). At P14, expression was mostly comparable to control mice although increased expression in the GCL was noted. BMP7 expression at P30 was clearly increased in Bmp7^ncko^ mice, particularly in the GCL, OPL and ONL. This dynamic change in expression was unexpected. Whereas in the Bmp7^ctrl^ retina BMP7 expression was reduced with the maturation of retina, expression in the Bmp7^ncko^ retina increased with maturation. Since BMP7 was strongly reduced in the Bmp7^ncko^ retina at P0, we investigated whether its loss affected the development of neurons and glial cells (Figure 6C). Much of the BMP7 appeared to associate with GFAP-positive glial cells, whereas no comparable colocalization was observed with NF-H (n=3 mice). An inverse relationship of BMP7 with NF-H expression was observed, whereas a direct relationship was found for GFAP when comparing Bmp7^ctrl^ and Bmp7^ncko^ retinas. This non-NCC-specific expression of BMP7 prompted us to query which retina cell types might be its source in the developing and adult retina.

### 3.8. BMP7 expression throughout retina development occurs in cells largely distinct from factors affected by its loss

Cell types expressing BMP7 and their relative location in the eye are summarized in Figure 7, top panel. Meta-analysis of scRNA datasets for the retina at various developmental stages (Clark et al., 2019) revealed the extent and cellular identity of BMP7-expressing cells. BMP7 (red) was observed predominantly in early and late retinal progenitor cells at embryonic and early post-natal stages. After birth, BMP7 was mainly localized to amacrine cells and some bipolar cells with minimal expression in retinal ganglion cells, cone and rod photoreceptors. Co-localization with markers affected by loss of BMP7 (CALB, NF-H, PAX6, RHO, S-OP and RCVRN) during embryonic retina development showed that expression of BMP7 overlapped with PAX6, and to some degree with CALB and NF-H. Co-expression with rod/cone markers was minimal (Figure 7, bottom panel).

**Figure 7.**
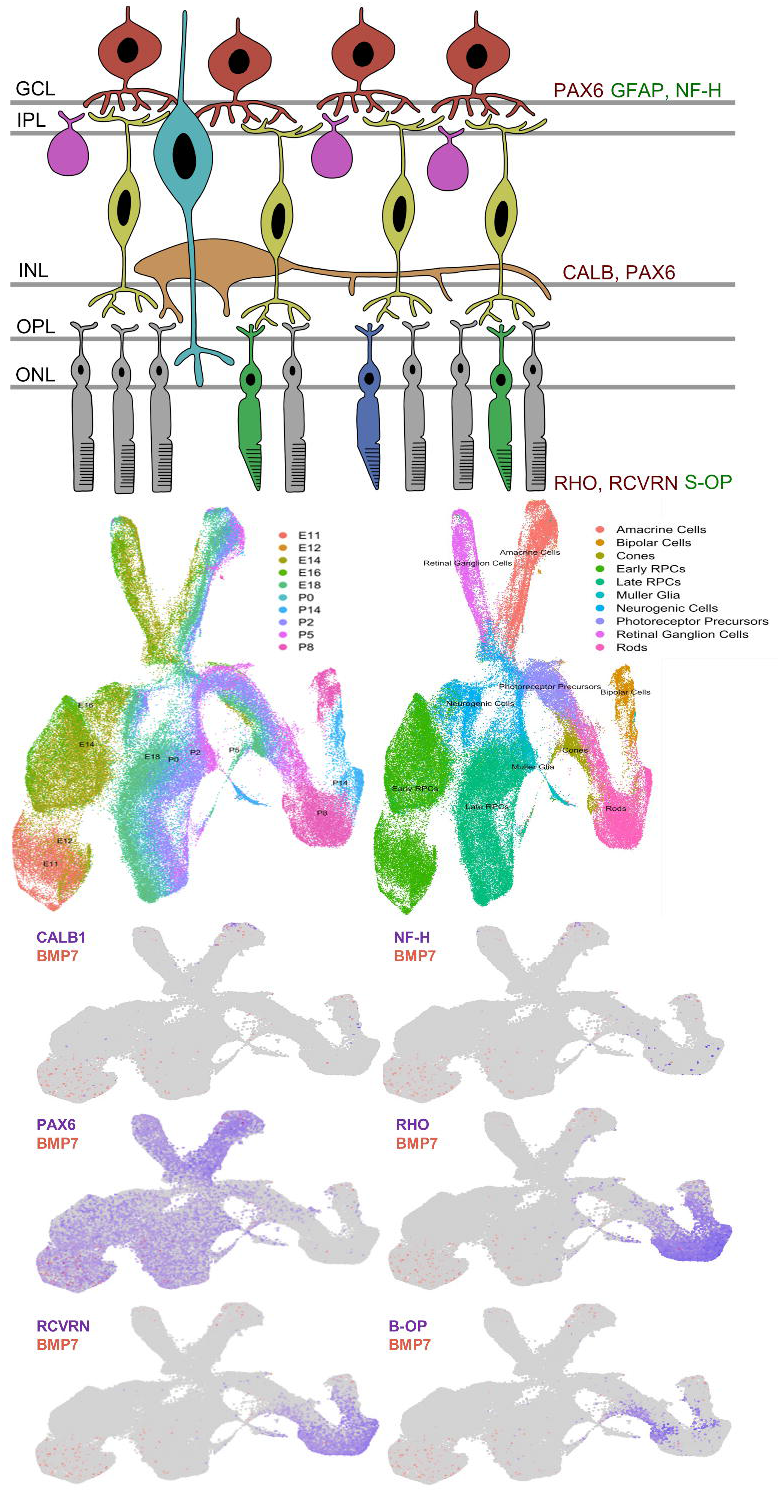
Co-expression of BMP7 with various proteins expressed by neurons and photoreceptors. **(Top panel)** A summary of aberrant proteins expression observed in P30 Bmp7^ncko^ mice retina. Increased expression of proteins in green and proteins with decreased expression in red. BMP7 expression in blue. **(Bottom panel)** Data mining of single cell RNA-Seq confirmed dynamic expression pattern of BMP7 during development along with co-expression of proteins (CALB, NF-H, PAX6, RHO, S-OP, RCVRN) assessed in this study. E: embryonic day; P: postnatal day. RHO: rhodopsin; RCVRN: recoverin; S-OP: blue opsin; CALB: calbindin; NF-H: neurofilament heavy; PAX6: paired box protein 6.

Overall, our data establish that BMP7 is an important trophic factor for retinal maturation and that NC-derived cells in the retina coordinate the development of different retinal cellular components.

## 4. Discussion

BMP7 has long been known as a critical factor for early eye development (Dudley and Robertson, 1997) In this study, we establish that BMP7 is expressed at postnatal stages of eye development and in the adult eye, some of which overlap with NC-derived structures. We show the functional importance of NC-derived BMP7 for the normal organization and function of the adult retina. This is to our knowledge the first report using a genetic approach to specifically address the role of BMP7 in a subset of cells in the adult eye. It is also the first study to demonstrate a requirement for an NC-derived growth factor for normal maturation and function of the retina.

BMP7 is expressed in the retina at various developmental stages based on the meta-analysis of scRNA datasets (Clark et al., 2019). BMP7 expression was observed in early and late retinal progenitor cells at embryonic and early post-natal stages. LacZ reporting of BMP7 supports this wider expression of BMP7. Although much of this expression is not confined to NC, in the postnatal eye, BMP7 is expressed in NC-derived structures of the cornea, the ciliary body and the iris. It should be noted that LacZ reporting identifies cells expressing the mRNA of the gene of interest while antibody staining demonstrates protein localization. Similar to what has been shown for the developing cortex (Segklia et al., 2012), the actual localization of BMP7 can be quite different from source of expression. BMP7 was recently shown to modulate corneal stroma and epithelial cell functions (Kowtharapu et al., 2018), we did not note any obvious changes to the cornea in Bmp7^ncko^ mice. BMP signaling is required for ciliary body development and both BMP4 and BMP7 have been implied in this process (Zhao et al., 2002) BMP4 loss-of-function studies failed to establish a direct role on ciliary body development (Rausch et al., 2018). In Bmp7^ncko^ mice, pupils appeared smaller, but the difference was not significant following their pharmacological dilation.

Based on persistence of NC-derived cells in the retina (Trost et al., 2016; Trost et al., 2016), their implication in ocular repair (Liu et al., 2014), and the importance of NC-derived BMP7 for neural stem cells and neurogenesis (Choe et al., 2013; Segklia et al., 2012), we focused our functional analysis on the retina. Changes to the anterior segment of the eye can affect functional analysis of the posterior segment, in particular ERG readings (Gagné et al., 2010; Johnson et al., 2019; Miura et al., 2016). Overall, the type and degree of differences observed in Bmp7^ncko^ mice for ERG measurements (a/b-wave, scotopic/photopic, flicker/implicit time) strongly suggest that they are not solely the consequence of changes in the anterior segment (Gagné et al., 2010; Johnson et al., 2019; Miura et al., 2016), but the result of specific changes to the inner and outer retina itself.

The reduced a-wave amplitude under scotopic conditions along with the reduction in rhodopsin and recoverin point towards impaired rod photoreceptor hyperpolarization (Figure 5) (Robson et al., 2003). Hyperpolarization can be affected if cytoskeleton-mediated alignment of photoreceptors is altered (Eckmiller, 2004). BMP7 stabilizes microtubules via the activation of c-Jun N-terminal kinases in neuronal cells (Podkowa et al., 2010); thus loss of BMP7 may similarly precipitate perturbed cone photoreceptor arrangement. Increased numbers of S-opsin expressing cone photoreceptors could be the consequence of a disorganized ONL. The a-wave amplitude reductions point to both rod and cone phototransduction defects, which in turn would contribute to b-wave amplitude reductions.

The b-wave amplitude and implicit time, reflecting inner retina function (Stockton and Slaughter, 1989), were reduced under both scotopic and photopic conditions in Bmp7^ncko^ mice. We observed a reduction in Calbindin, indicating changes to bipolar, amacrine and horizontal cells, in agreement with our observed b-wave deficit. Overexpression of the BMP antagonist Noggin1 results in a reduction of bipolar cells in the INL (Messina et al., 2016). However, these changes appeared to be more qualitative than quantitative.

The reduced b/a amplitude ratios point to additional dysfunctions in the inner retina. Loss of NC-derived BMP7 led to increased retinal expression of BMP7 at P30. BMP7 has been associated with reactive gliosis and neuro-inflammation (Dharmarajan et al., 2017, 2014). Reactive gliosis is a universal reaction to neuronal injury (Buffo et al., 2008), typically resulting in proliferative gliosis (Vázquez-Chona et al., 2011). In contrast, hypertrophic gliosis is characterized by Müller cells getting larger while not sending projections in the outer retina (de Melo et al., 2012). The increase in Iba1 in P30 Bmp7^ncko^ retinas is in line with the development of a gliosis. The altered distribution of GFAP and NF-H again points towards involvement of BMP7 in cytoskeletal organization of glia and astrocytes. The reduction in PAX6 at P30 further indicates an effect on amacrine and RGCs cells. This may be similar to what has been described in the developing cortex, where loss of BMP7 affects radial glia progenitor cells (Segklia et al., 2012). On the other hand, increased expression of GFAP is associated with de-differentiation of Müller glia and regeneration of rod photoreceptors (Raymond et al., 2006), and thus might reflect retina stress caused by impaired rod photoreceptor function and disrupted neuronal circuitry (Lewis and Fisher, 2003; Raymond et al., 2006). It could be that NC-derived BMP7 provides trophic support to various neuroepithelial cells, similar to what has been described for the brain (Choe et al., 2013; Segklia et al., 2012). The unexpected dynamic expression of BMP7 (variable reduction at P0, comparable expression at P14 and particularly its increased expression in the P30 mutant retina) underscores the importance of BMP7 for the adult retina.

Our data reveals an important role for NCC-derived BMP7 in the developing retina. Lineage tracing was in line with previously described observations (Liu et al., 2014; Trost et al., 2013). In particular, using both Sox10-Cre and Wnt1-Cre reporter mice Liu et. al (Liu et al., 2014) suggested the existence of cells spanning the retina reminiscent of glia cells, which were observed in some but not all of our experiments (Figure 6A). There is controversy regarding the use of Wnt1-Cre to delete NCC, as ectopic expression of WNT1 has been observed in the ventral midbrain of this mouse (Lewis et al., 2013). A Wnt1-Cre2 mouse was developed to overcome this limitation (Lewis et al., 2013); however, this mouse was not used in this study due to its still insufficient characterization and emerging controversy on its faithfulness (Debbache et al., 2018). While the Wnt1-cre mouse has been shown to identify mostly neural crest cells, it cannot be formally excluded that Wnt1 may be expressed in a few non-neural crest cells at some stage during retina development. To gauge if the ERG phenotype observed in Bmp7^ncko^ mice could be the direct result of ectopic Wnt1-Cre activation, we tested heterozygote Bmp7^ncko^ mice. Their ERG responses were comparable to Bmp7^ctrl^ mice; hence, we have no reason to believe that phenotypic observations in the Bmp7^ncko^ mice are a direct consequence of ectopic Wnt1 expression.

Loss of function mutations of BMP7 in humans are associated with ocular defects, although no specific details on retina function were described (Wyatt et al., 2010). At the same time, these patients showed various craniofacial defects overlapping in spectrum with Stickler syndrome. Bmp7^ncko^ mice also display craniofacial defects reminiscent of Stickler syndrome (Baddam et al., 2021; Kouskoura et al., 2013). Alterations to BMP signaling have been associated with Stickler syndrome (Nixon et al., 2019), and the genes mutated in Stickler syndrome (Collagens II, IX, XI) are expressed at reduced levels in BMP7-mutant embryos (data not shown). Stickler syndrome patients often display reduced scotopic b-wave amplitudes (Kondo et al., 2020) similar to Bmp7^ncko^ mice. However, they demonstrate prolonged b-wave implicit times in contrast to Bmp7^ncko^ mice that show faster b-wave implicit times. Beyond those associated with high myopia, the visual acuity deficits in Sticker Syndrome patients are not understood; the cellular and molecular alterations underlying the deficits remain to be characterized. Molecular findings from this study may provide clues to better understand the functional deficits.

In summary, this study demonstrates for the first time that NCC, in part mediated through secretion of growth factors such as BMP7, contributes to cell organization and neuronal circuitry in the adult retina. The observed phenotype is likely the consequence of early alterations to retina progenitors that manifest as visual function deficits post-development, but also indicates a continued requirement for BMP7 in the adult retina. Further studies are needed to unravel in more detail how neural crest contributes to retina development and maturation.

## 5. Acknowledgements

The authors would like to acknowledge Luc Berthiaume (Department of Cell Biology, University of Alberta) for kindly providing the anti-GFP antibody used in this study and Jennifer Hocking (Department of Surgery, University of Alberta) and Pierre Mattar (Cellular and Molecular Medicine, University of Ottawa) for critical reading of the manuscript.

## References

Baddam, P., Biancardi, V., Roth, D.M., Eaton, F., Thereza-Bussolaro, C., Mandal, R., Wishart, D.S., Barr, A., MacLean, J., Flores-Mir, C., Pagliardini, S., Graf, D., 2021. Neural crest-specific deletion of Bmp7 leads to midfacial hypoplasia, nasal airway obstruction, and disordered breathing modelling Obstructive Sleep Apnea. Dis. Model. Mech. dmm.047738. https://doi.org/10.1242/dmm.047738

Baddam, P., Kung, T., Adesida, A.B., Graf, D., 2020. Histological and molecular characterization of the growing nasal septum in mice. J Anat joa.13332. https://doi.org/10.1111/joa.13332

Bassett, E.A., Wallace, V.A., 2012. Cell fate determination in the vertebrate retina. Trends in Neurosciences 35, 565–573. https://doi.org/10.1016/j.tins.2012.05.004

Buffo, A., Rite, I., Tripathi, P., Lepier, A., Colak, D., Horn, A.-P., Mori, T., Gotz, M., 2008. Origin and progeny of reactive gliosis: A source of multipotent cells in the injured brain. Proceedings of the National Academy of Sciences 105, 3581–3586. https://doi.org/10.1073/pnas.0709002105

Cheng, N., Pagtalunan, E., Abushaibah, A., Naidu, J., Stell, W.K., Rho, J.M., Sauvé, Y., 2020. Atypical visual processing in a mouse model of autism. Sci Rep 10, 12390. https://doi.org/10.1038/s41598-020-68589-9

Choe, Y., Kozlova, A., Graf, D., Pleasure, S.J., 2013. Bone Morphogenic Protein Signaling Is a Major Determinant of Dentate Development. Journal of Neuroscience 33, 6766–6775. https://doi.org/10.1523/JNEUROSCI.0128-13.2013

Clark, B.S., Stein-O’Brien, G.L., Shiau, F., Cannon, G.H., Davis-Marcisak, E., Sherman, T., Santiago, C.P., Hoang, T.V., Rajaii, F., James-Esposito, R.E., Gronostajski, R.M., Fertig, E.J., Goff, L.A., Blackshaw, S., 2019. Single-Cell RNA-Seq Analysis of Retinal Development Identifies NFI Factors as Regulating Mitotic Exit and Late-Born Cell Specification. Neuron 102, 1111-1126.e5. https://doi.org/10.1016/j.neuron.2019.04.010

Creel, D.J., 1995. Clinical Electrophysiology, in: Kolb, H., Fernandez, E., Nelson, R. (Eds.), Webvision: The Organization of the Retina and Visual System. University of Utah Health Sciences Center, Salt Lake City (UT).

de Melo, J., Miki, K., Rattner, A., Smallwood, P., Zibetti, C., Hirokawa, K., Monuki, E.S., Campochiaro, P.A., Blackshaw, S., 2012. Injury-independent induction of reactive gliosis in retina by loss of function of the LIM homeodomain transcription factor Lhx2. Proceedings of the National Academy of Sciences 109, 4657–4662. https://doi.org/10.1073/pnas.1107488109

Debbache, J., Parfejevs, V., Sommer, L., 2018. Cre-driver lines used for genetic fate mapping of neural crest cells in the mouse: An overview. genesis 56, e23105. https://doi.org/10.1002/dvg.23105

Dharmarajan, S., Fisk, D.L., Sorenson, C.M., Sheibani, N., Belecky-Adams, T.L., 2017. Microglia activation is essential for BMP7-mediated retinal reactive gliosis. J Neuroinflammation 14, 76. https://doi.org/10.1186/s12974-017-0855-0

Dharmarajan, S., Gurel, Z., Wang, S., Sorenson, C.M., Sheibani, N., Belecky-Adams, T.L., 2014. Bone morphogenetic protein 7 regulates reactive gliosis in retinal astrocytes and Müller glia. Mol Vis 20, 1085–1108.

Dudley, A.T., Robertson, E.J., 1997. Overlapping expression domains of bone morphogenetic protein family members potentially account for limited tissue defects in BMP7 deficient embryos. Dev Dyn 208, 349–362. https://doi.org/10.1002/(SICI)1097-0177(199703)208:3<349::AID-AJA6>3.0.CO;2-I

Eckmiller, M., 2004. Defective cone photoreceptor cytoskeleton, alignment, feedback, and energetics can lead to energy depletion in macular degeneration. Progress in Retinal and Eye Research 23, 495–522. https://doi.org/10.1016/j.preteyeres.2004.04.005

Ellis R. Hematoxylin and Eosin (H&E) Staining Protocol. In: IHC World. http://www.ihcworld.com/_protocols/special_stains/h&e_ellis.htm.

Etchevers, H.C., Vincent, C., Le Douarin, N.M., Couly, G.F., 2001. The cephalic neural crest provides pericytes and smooth muscle cells to all blood vessels of the face and forebrain. Development 128, 1059–1068.

Gage, P.J., Rhoades, W., Prucka, S.K., Hjalt, T., 2005. Fate maps of neural crest and mesoderm in the mammalian eye. Invest Ophthalmol Vis Sci 46, 4200–4208. https://doi.org/10.1167/iovs.05-0691

Gagné, A.-M., Lavoie, J., Lavoie, M.-P., Sasseville, A., Charron, M.-C., Hébert, M., 2010. Assessing the impact of non-dilating the eye on full-field electroretinogram and standard flash response. Doc Ophthalmol 121, 167–175. https://doi.org/10.1007/s10633-010-9242-1

Godin, R.E., Takaesu, N.T., Robertson, E.J., Dudley, A.T., 1998. Regulation of BMP7 expression during kidney development. Development 125, 3473.

Gu, Y.-N., Lee, E.-S., Jeon, C.-J., 2016. Types and density of calbindin D28k-immunoreactive ganglion cells in mouse retina. Experimental Eye Research 145, 327–336. https://doi.org/10.1016/j.exer.2016.02.001

Hoang, T., Wang, J., Boyd, P., Wang, F., Santiago, C., Jiang, L., Yoo, S., Lahne, M., Todd, L.J., Jia, M., Saez, C., Keuthan, C., Palazzo, I., Squires, N., Campbell, W.A., Rajaii, F., Parayil, T., Trinh, V., Kim, D.W., Wang, G., Campbell, L.J., Ash, J., Fischer, A.J., Hyde, D.R., Qian, J., Blackshaw, S., 2020. Gene regulatory networks controlling vertebrate retinal regeneration. Science 370, eabb8598. https://doi.org/10.1126/science.abb8598

Jeon, C.-J., Strettoi, E., Masland, R.H., 1998. The Major Cell Populations of the Mouse Retina. J. Neurosci. 18, 8936–8946. https://doi.org/10.1523/JNEUROSCI.18-21-08936.1998

Johnson, M.A., Jeffrey, B.G., Messias, A.M.V., Robson, A.G., 2019. ISCEV extended protocol for the stimulus–response series for the dark-adapted full-field ERG b-wave. Doc Ophthalmol 138, 217–227. https://doi.org/10.1007/s10633-019-09687-6

Kashiwagi, K., Ou, B., Nakamura, S., Tanaka, Y., Suzuki, M., Tsukahara, S., 2003. Increase in Dephosphorylation of the Heavy Neurofilament Subunit in the Monkey Chronic Glaucoma Model. Invest. Ophthalmol. Vis. Sci. 44, 154. https://doi.org/10.1167/iovs.02-0398

Kondo, H., Fujimoto, K., Imagawa, M., Oku, K., Matsushita, I., Hayashi, T., Nagata, T., 2020. Electroretinograms of eyes with Stickler syndrome. Doc Ophthalmol 140, 233–243. https://doi.org/10.1007/s10633-019-09739-x

Kouskoura, T., Kozlova, A., Alexiou, M., Blumer, S., Zouvelou, V., Katsaros, C., Chiquet, M., Mitsiadis, T.A., Graf, D., 2013. The Etiology of Cleft Palate Formation in BMP7-Deficient Mice. PLoS ONE 8, e59463. https://doi.org/10.1371/journal.pone.0059463

Kowtharapu, B., Prakasam, R., Murín, R., Koczan, D., Stahnke, T., Wree, A., Jünemann, A., Stachs, O., 2018. Role of Bone Morphogenetic Protein 7 (BMP7) in the Modulation of Corneal Stromal and Epithelial Cell Functions. IJMS 19, 1415. https://doi.org/10.3390/ijms19051415

Langenberg, T., Kahana, A., Wszalek, J.A., Halloran, M.C., 2008. The eye organizes neural crest cell migration. Dev. Dyn. 237, 1645–1652. https://doi.org/10.1002/dvdy.21577

Lewis, A.E., Vasudevan, H.N., O’Neill, A.K., Soriano, P., Bush, J.O., 2013. The widely used Wnt1-Cre transgene causes developmental phenotypes by ectopic activation of Wnt signaling. Developmental Biology 379, 229–234. https://doi.org/10.1016/j.ydbio.2013.04.026

Lewis, G.P., Fisher, S.K., 2003. Up-Regulation of Glial Fibrillary Acidic Protein in Response to Retinal Injury: Its Potential Role in Glial Remodeling and a Comparison to Vimentin Expression, in: International Review of Cytology. Elsevier, pp. 263–290. https://doi.org/10.1016/S0074-7696(03)30005-1

Li, S., Goldowitz, D., Swanson, D.J., 2007. The Requirement of Pax6 for Postnatal Eye Development: Evidence from Experimental Mouse Chimeras. Invest. Ophthalmol. Vis. Sci. 48, 3292. https://doi.org/10.1167/iovs.06-1482

Liu, B., Hunter, D.J., Smith, A.A., Chen, S., Helms, J.A., 2014. The capacity of neural crest-derived stem cells for ocular repair: Neural Crest-Derived Stem Cells for Ocular Repair. Birth Defect Res C 102, 299–308. https://doi.org/10.1002/bdrc.21077

Malik, Z., Roth, D.M., Eaton, F., Theodor, J.M., Graf, D., 2020. Mesenchymal Bmp7 Controls Onset of Tooth Mineralization: A Novel Way to Regulate Molar Cusp Shape. Front. Physiol. 11, 698. https://doi.org/10.3389/fphys.2020.00698

Martins, R.R., Zamzam, M., Moosajee, M., Thummel, R., Henriques, C.M., MacDonald, R.B., 2020. Müller Glia regenerative potential is maintained throughout life despite neurodegeneration and gliosis in the ageing zebrafish retina (preprint). Neuroscience. https://doi.org/10.1101/2020.06.28.174821

Masland, R.H., 2012. The neuronal organization of the retina. Neuron 76, 266–280. https://doi.org/10.1016/j.neuron.2012.10.002

McCabe, J.B., Berthiaume, L.G., 1999. Functional Roles for Fatty Acylated Amino-terminal Domains in Subcellular Localization. MBoC 10, 3771–3786. https://doi.org/10.1091/mbc.10.11.3771

Messina, A., Bridi, S., Bozza, A., Bozzi, Y., Baudet, M.-L., Casarosa, S., 2016. Noggin 1 overexpression in retinal progenitors affects bipolar cell generation. Int. J. Dev. Biol. 60, 151–157. https://doi.org/10.1387/ijdb.150402am

Miura, G., Nakamura, Y., Sato, E., Yamamoto, S., 2016. Effects of cataracts on flicker electroretinograms recorded with RETeval™ system: new mydriasis-free ERG device. BMC Ophthalmol 16, 22. https://doi.org/10.1186/s12886-016-0200-x

Morona, R., Moreno, N., Lopez, J.M., Muñoz, M., Domínguez, L., González, A., 2008. Calbindin-D28k and calretinin as markers of retinal neurons in the anuran amphibian Rana perezi. Brain Res Bull 75, 379–383. https://doi.org/10.1016/j.brainresbull.2007.10.026

Muzumdar, M.D., Tasic, B., Miyamichi, K., Li, L., Luo, L., 2007. A global double-fluorescent Cre reporter mouse. Genesis 45, 593–605. https://doi.org/10.1002/dvg.20335

Nadal-Nicolás, F.M., Kunze, V.P., Ball, J.M., Peng, B.T., Krishnan, A., Zhou, G., Dong, L., Li, W., 2020. True S-cones are concentrated in the ventral mouse retina and wired for color detection in the upper visual field. eLife 9, e56840. https://doi.org/10.7554/eLife.56840

Nixon, T.R.W., Richards, A., Towns, L.K., Fuller, G., Abbs, S., Alexander, P., McNinch, A., Sandford, R.N., Snead, M.P., 2019. Bone morphogenetic protein 4 (BMP4) loss-of-function variant associated with autosomal dominant Stickler syndrome and renal dysplasia. Eur J Hum Genet 27, 369–377. https://doi.org/10.1038/s41431-018-0316-y

Ortín-Martínez, A., Nadal-Nicolás, F.M., Jiménez-López, M., Alburquerque-Béjar, J.J., Nieto-López, L., García-Ayuso, D., Villegas-Pérez, M.P., Vidal-Sanz, M., Agudo-Barriuso, M., 2014. Number and Distribution of Mouse Retinal Cone Photoreceptors: Differences between an Albino (Swiss) and a Pigmented (C57/BL6) Strain. PLOS ONE 9, e102392. https://doi.org/10.1371/journal.pone.0102392

Podkowa, M., Zhao, X., Chow, C.-W., Coffey, E.T., Davis, R.J., Attisano, L., 2010. Microtubule Stabilization by Bone Morphogenetic Protein Receptor-Mediated Scaffolding of c-Jun N-Terminal Kinase Promotes Dendrite Formation. MCB 30, 2241–2250. https://doi.org/10.1128/MCB.01166-09

Rausch, R.L., Libby, R.T., Kiernan, A.E., 2018. Ciliary margin-derived BMP4 does not have a major role in ocular development. PLoS ONE 13, e0197048. https://doi.org/10.1371/journal.pone.0197048

Raymond, P.A., Barthel, L.K., Bernardos, R.L., Perkowski, J.J., 2006. Molecular characterization of retinal stem cells and their niches in adult zebrafish. BMC Dev Biol 6, 36. https://doi.org/10.1186/1471-213X-6-36

Reichenbach, A., Bringmann, A., 2013. New functions of Müller cells: New Functions of Müller Cells. Glia 61, 651–678. https://doi.org/10.1002/glia.22477

Richmond R. Davidson’s Fixative Protocol. In: IHC World. http://www.ihcworld.com/_protocols/histology/davidson_fixative.htm.

Riesenberg, A.N., Le, T.T., Willardsen, M.I., Blackburn, D.C., Vetter, M.L., Brown, N.L., 2009. Pax6 regulation of Math5 during mouse retinal neurogenesis. genesis 47, 175–187. https://doi.org/10.1002/dvg.20479

Robson, J.G., Saszik, S.M., Ahmed, J., Frishman, L.J., 2003. Rod and cone contributions to the a -wave of the electroretinogram of the macaque. The Journal of Physiology 547, 509–530. https://doi.org/10.1113/jphysiol.2002.030304

Sarthy, P.V., Fu, M., Huang, J., 1991. Developmental expression of the Glial fibrillary acidic protein (GFAP) gene in the mouse retina. Cell Mol Neurobiol 11, 623–637. https://doi.org/10.1007/BF00741450

Segklia, A., Seuntjens, E., Elkouris, M., Tsalavos, S., Stappers, E., Mitsiadis, T.A., Huylebroeck, D., Remboutsika, E., Graf, D., 2012. Bmp7 Regulates the Survival, Proliferation, and Neurogenic Properties of Neural Progenitor Cells during Corticogenesis in the Mouse. PLoS ONE 7, e34088. https://doi.org/10.1371/journal.pone.0034088

Shen, J., Colonnese, M.T., 2016. Development of Activity in the Mouse Visual Cortex. Journal of Neuroscience 36, 12259–12275. https://doi.org/10.1523/JNEUROSCI.1903-16.2016

Siegenthaler, J.A., Pleasure, S.J., 2011. We have got you ‘covered’: how the meninges control brain development. Current Opinion in Genetics & Development 21, 249–255. https://doi.org/10.1016/j.gde.2010.12.005

Stockton, R.A., Slaughter, M.M., 1989. B-wave of the electroretinogram. A reflection of ON bipolar cell activity. Journal of General Physiology 93, 101–122. https://doi.org/10.1085/jgp.93.1.101

Tassoni, A., Gutteridge, A., Barber, A.C., Osborne, A., Martin, K.R., 2015. Molecular Mechanisms Mediating Retinal Reactive Gliosis Following Bone Marrow Mesenchymal Stem Cell Transplantation: Retinal Glial Responses to Transplanted MSCs. Stem Cells 33, 3006–3016. https://doi.org/10.1002/stem.2095

Tkatchenko, T.V., Shen, Y., Tkatchenko, A.V., 2010. Analysis of Postnatal Eye Development in the Mouse with High-Resolution Small Animal Magnetic Resonance Imaging. Invest. Ophthalmol. Vis. Sci. 51, 21. https://doi.org/10.1167/iovs.08-2767

Trost, A., Lange, S., Schroedl, F., Bruckner, D., Motloch, K.A., Bogner, B., Kaser-Eichberger, A., Strohmaier, C., Runge, C., Aigner, L., Rivera, F.J., Reitsamer, H.A., 2016. Brain and Retinal Pericytes: Origin, Function and Role. Front. Cell. Neurosci. 10. https://doi.org/10.3389/fncel.2016.00020

Trost, A., Schroedl, F., Lange, S., Rivera, F.J., Tempfer, H., Korntner, S., Stolt, C.C., Wegner, M., Bogner, B., Kaser-Eichberger, A., Krefft, K., Runge, C., Aigner, L., Reitsamer, H.A., 2013. Neural Crest Origin of Retinal and Choroidal Pericytes. Invest. Ophthalmol. Vis. Sci. 54, 7910. https://doi.org/10.1167/iovs.13-12946

Vázquez-Chona, F.R., Swan, A., Ferrell, W.D., Jiang, L., Baehr, W., Chien, W.-M., Fero, M., Marc, R.E., Levine, E.M., 2011. Proliferative reactive gliosis is compatible with glial metabolic support and neuronal function. BMC Neurosci 12, 98. https://doi.org/10.1186/1471-2202-12-98

Verma, R., Pianta, M.J., 2009. The contribution of human cone photoreceptorsto the photopic flicker electroretinogram. Journal of Vision 9, 9–9. https://doi.org/10.1167/9.3.9

Wawersik, S., Purcell, P., Rauchman, M., Dudley, A.T., Robertson, E.J., Maas, R., 1999. BMP7 acts in murine lens placode development. Dev Biol 207, 176–188. https://doi.org/10.1006/dbio.1998.9153

Whikehart, D.R., 2010. Corneal Endothelium: Overview, in: Encyclopedia of the Eye. Elsevier, pp. 424–434. https://doi.org/10.1016/B978-0-12-374203-2.00074-9

Williams, A.L., Bohnsack, B.L., 2015. Neural crest derivatives in ocular development: Discerning the eye of the storm: Neural Crest Derivatives in Eye Development. Birth Defect Res C 105, 87–95. https://doi.org/10.1002/bdrc.21095

Wyatt, A.W., Osborne, R.J., Stewart, H., Ragge, N.K., 2010. Bone morphogenetic protein 7 (BMP7) mutations are associated with variable ocular, brain, ear, palate, and skeletal anomalies. Hum. Mutat. 31, 781–787. https://doi.org/10.1002/humu.21280

Zhao, S., Chen, Q., Hung, F.-C., Overbeek, P.A., 2002. BMP signaling is required for development of the ciliary body. Development 129, 4435–4442.

Zouvelou, V., Luder, H.-U., Mitsiadis, T.A., Graf, D., 2009. Deletion of BMP7 affects the development of bones, teeth, and other ectodermal appendages of the orofacial complex. J. Exp. Zool. 312B, 361–374. https://doi.org/10.1002/jez.b.21262

