## Supplemental Figures for "Wnt1-Cre mediated deletion of BMP7 suggests a role for neural crest-derived BMP7 in retina development and function"

**Supplementary Figures**

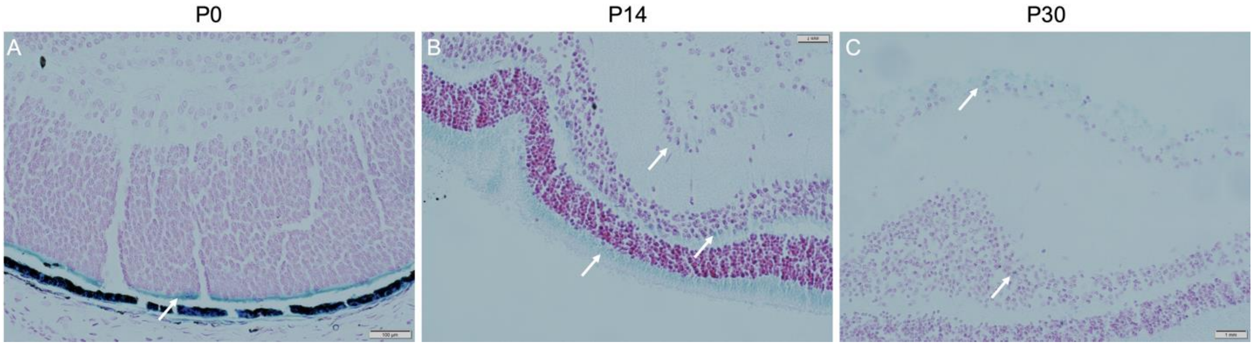


**Supplemental Figure 1: Higher magnification images of Bmp7 expression.** 40X images of **(A)** P0, **(B)** P14, **(C)** P30 Bmp7LacZ retina. P0: postnatal day 0; P14: postnatal day 14; P30: postnatal day 30.

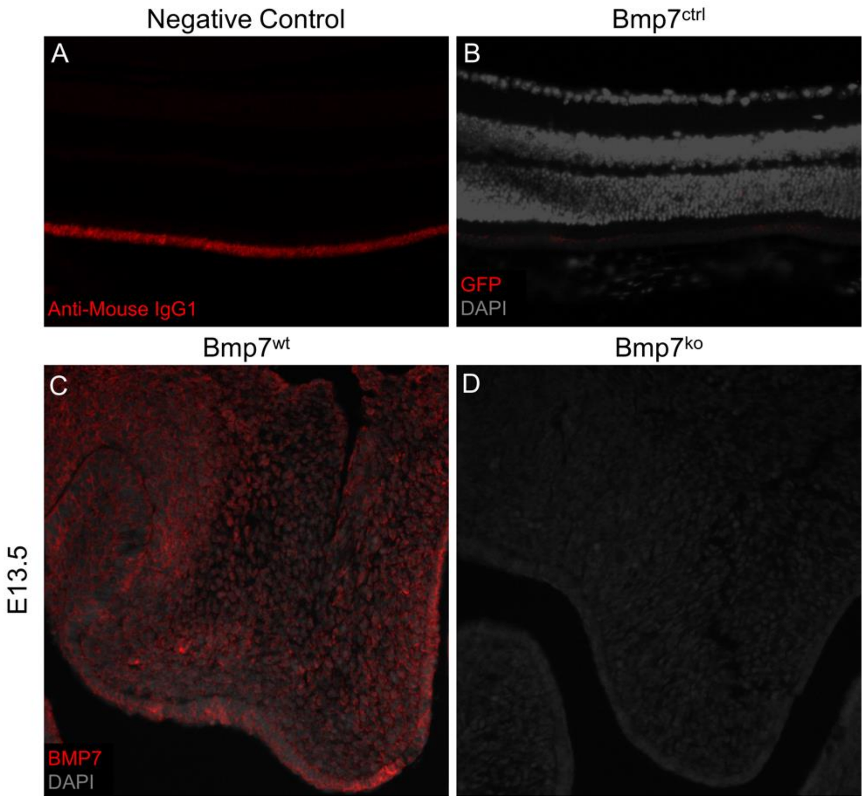


**Supplemental Figure 2: Immunofluorescence antibody controls.** Negative control **(A)** of Anti-Mouse IgG1 AF647 antibody demonstrates background staining in RPE. Negative GFP staining for neural crest cells observed in P30 Bmp7$^{ctrl}$ mice **(B)**. Specificity of Bmp7 antibody was tested using paraffin sections of palatal shelves in E13.5 control (Bmp7$^{wt}$) **(C)** and Bmp7 knockout (Bmp7$^{ko}$) mice **(D)**. P30: postnatal day 30; RPE: retinal pigmented epithelium; gfp: green fluorescent protein; Bmp7: bone morphogenetic protein 7; e13.5: embryonic day 13.5.

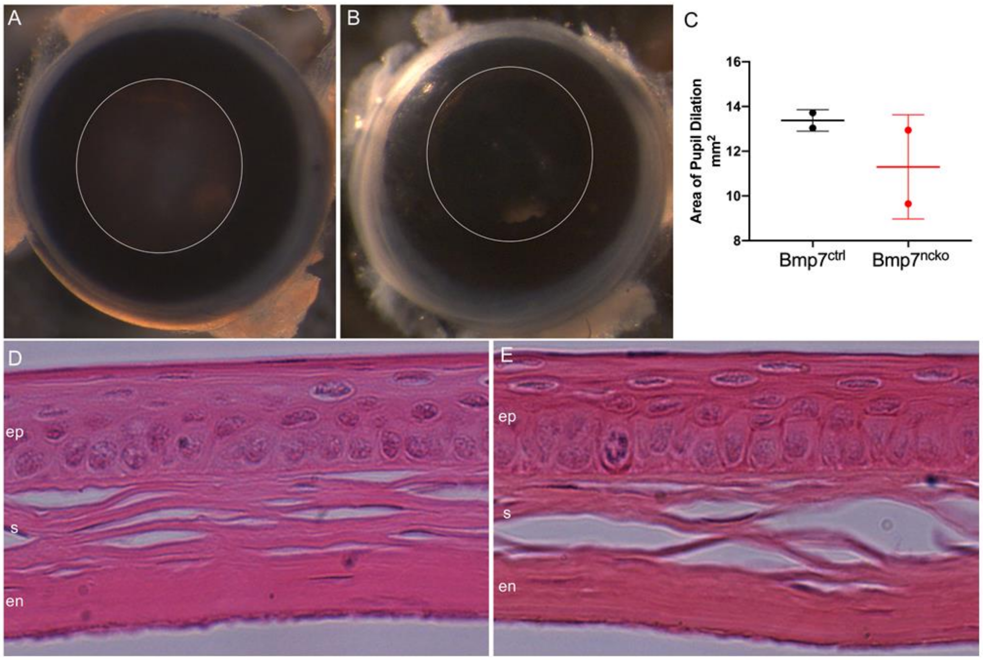


**Supplemental Figure 3: No obvious differences in anterior structures of P30 Bmp7$^{ncko}$ retina.** Bmp7$^{ctrl}$ **(A)** and Bmp7$^{ncko}$ **(B)** pupils after pharmacological dilation demonstrating no difference in degree of pupil dilation **(C)**. No structural differences observed in P30 Bmp7$^{ctrl}$ **(D)** and Bmp7$^{ncko}$ **(E)** cornea. ep: epithelium; s: stroma; en: endothelium. (n=3/genotype). P30: postnatal day 30.

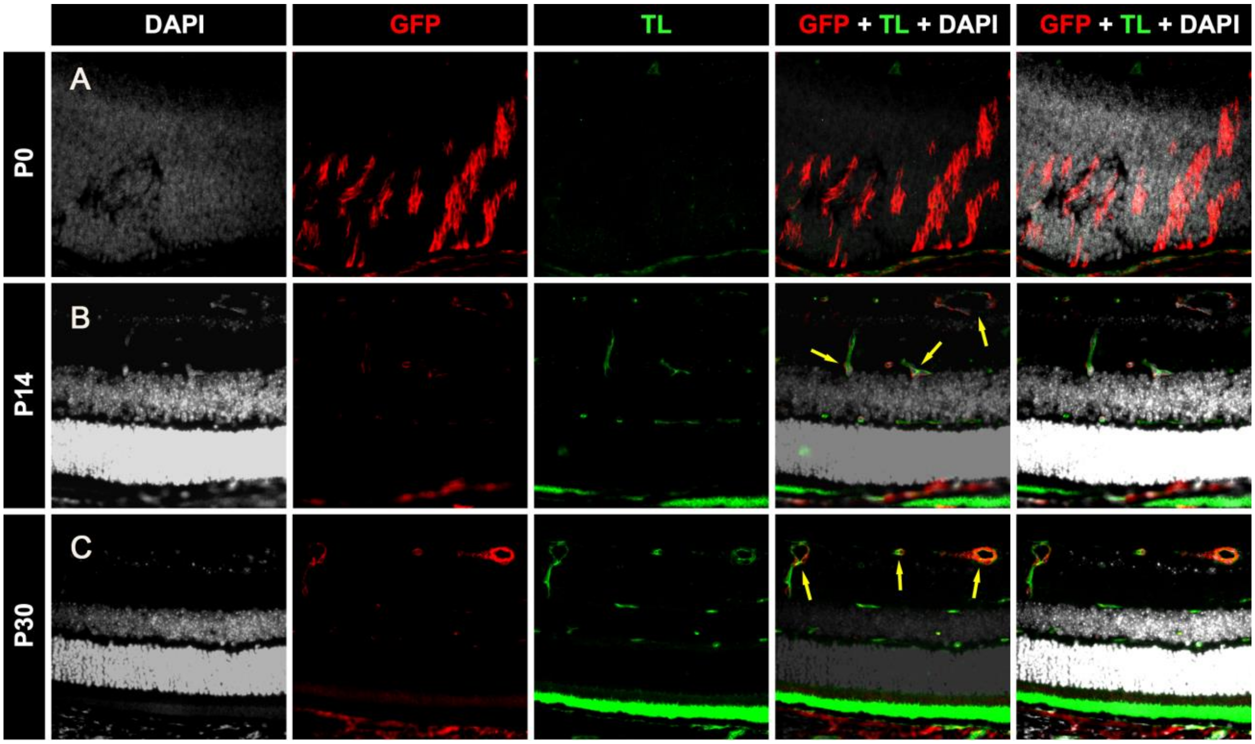


**Supplemental Figure 4: Higher magnification images of expression.** 40X images of P0, **(B)** P14, **(C)** P30 Bmp7LacZ retina. Bmp7 knockout ($Bmp7^{ko}$) mice. P0: postnatal day 0; P14: postnatal day 14; P30: postnatal day 30. Yellow arrows denote co-localized cells expressing both GFP and TL.
